# Stochastic simulations show how passive immunization can influence the germinal centre reaction and optimize host humoral responses

**DOI:** 10.1101/441691

**Authors:** Amar K. Garg, Rajat Desikan, Narendra M. Dixit

**Author notes:** Shared first authorship (Amar K. Garg, Rajat Desikan).

## Abstract

Passive immunization with antigen-specific antibodies was shown recently to induce lasting improvements in endogenous antibody production, raising the prospect of using passive immunization as a tool to engineer host humoral responses. The mechanism with which administered antibodies alter endogenous antibody production remains unknown. B cells that produce antigen-specific antibodies evolve and get selected in germinal centres (GCs). This selection requires that B cells acquire antigen presented in GCs. We hypothesized that passive immunization biases this selection in favour of B cells with high affinities for antigen. Administered antibodies form immune complexes with antigen which only B cells with higher affinities than the administered antibodies for antigen can rupture and acquire antigen, thus increasing the selection stringency in GCs. With this mechanistic hypothesis, we constructed a stochastic simulation model of the GC reaction. The simulations recapitulated and synthesized several independent experimental observations, presenting strong evidence in support of our hypothesis. Further, the simulations revealed a quality-quantity trade-off constraining the GC response. As the selection stringency increased, surviving B cells had higher affinities for antigen but fewer B cells survived. Increasing antigen availability in the GC relaxed this constraint. The affinity of the administered antibodies and/or antigen availability could thus be tuned to maximize the GC output. Comprehensively spanning parameter space, we predict passive immunization protocols that exploit the quality-quantity trade-off and maximize the GC output. Our study thus presents a new conceptual understanding of the GC reaction and a computational framework for the rational optimization of passive immunization strategies.

**Significance statement:** When natural antibody production is inadequate, passive immunization with external antibodies can alleviate disease. Remarkably, passive immunization induced lasting improvements in natural antibody production in recent studies, suggesting that it could be deployed to engineer natural antibody responses. However, how administered antibodies alter natural antibody production remains unknown. B cells that produce antibodies targeting specific antigen evolve in germinal centres (GCs). We hypothesized that administered antibodies form complexes with antigen, preferentially allowing B cells with higher affinities to acquire antigen and be selected, thus altering antibody production. With this mechanistic hypothesis, we performed stochastic simulations of the GC reaction, which recapitulated experiments, unravelled a quality-quantity trade-off constraining the GC response, and predicted passive immunization protocols that maximized the GC output.

## Introduction

The canonical mechanism of disease control by passive immunization with pre-formed antibodies (Abs) is the immediate neutralization and clearance of antigen (Ag).^1–4^ This mechanism underlies the array of Ab therapeutics against pathogens such as HIV-1^3,5^, influenza^3,6,7^, and respiratory syncytial virus (RSV)^3,8^, and against auto-immune disorders^9^ and cancer^2,10^. It is also what results in the acquisition of immunity by infants from mothers by the transfer of Abs through the placenta or breast milk^2^. Transcending this canonical mechanism, recent studies have shown that passive immunization with Ag-specific Abs can modulate the evolution of host B cell responses to the Ag.^11–15^ For instance, HIV-1 infected individuals infused with a single dose of the broadly neutralizing antibody (bNAb) 3BNC117 developed endogenous serum Ab responses with significantly improved breadth and potency compared to untreated individuals.^15^ The improvement was observed well after the administered bNAbs were cleared from circulation. Similarly, passive administration of low dose neutralizing Abs to newborn macaques before SHIV challenge improved the production of endogenous neutralizing Abs, the presence of which correlated with low set-point viremia and 100% survival.^13^ These studies suggest that passive immunization protocols that elicit potent, lasting humoral responses can be devised. A key limitation is the poor understanding of the mechanism with which passively administered Abs alter the humoral response. Passive immunization is a drug-like therapy with exogenous Abs targeting specific Ag.^2^ How it influences endogenous Ab production is puzzling. Here, we propose a mechanism with which passive immunization can influence endogenous Ab production and can therefore be exploited as a tool to engineer the humoral response.

Abs are produced in germinal centres (GCs; Figure 1*a*), which are temporary structures formed in lymphoid organs during an infection, where B cells that can produce Abs with high affinity for a target Ag evolve and get selected.^16–18^ GC B cells exist in a default pro-apoptotic state and must receive two signals to survive and proliferate^16,17^: Ag uptake from follicular dendritic cells (FDCs) and, subsequently, help from follicular T helper (T_*fh*_) cells, both in the ‘light zone’ of the GC. Each B cell expresses a single type of B cell receptor (BCR).^16^ BCRs acquire Ag from immune complexes (ICs) attached to FC receptors and presented on FDCs.^18,19^ ICs are non-covalently bound complexes between Ag and Abs or complement fragments. The higher is the affinity of a BCR for Ag, the greater is the likelihood of the BCR acquiring Ag.^20^ Upon Ag acquisition, B cells internalize the Ag and present it as peptides loaded on major histocompatibility complex class II molecules (pMHCII) on their surfaces in amounts proportional to the quantum of acquired Ag.^16^ T_*fh*_ cells preferentially synapse with B cells displaying relatively high amounts of pMHCII, leading to their selection.^16^ Most B cells thus selected traffic to the ‘dark zone’ of the GC, where they proliferate and mutate.^16,17^ Mutations can alter the BCRs and hence their affinity for Ag. From the dark zone, B cells re-enter the light zone and are subjected to the same selection process again.^16,17,21^ B cells with BCRs of increasingly higher affinities for Ag are thus selected, a process termed affinity maturation (AM). A small minority of B cells selected exits the GC and differentiates into plasma and memory B cells.^16,22^ Mature plasma cells produce Abs in the serum and mount the humoral response.

**Figure 1.**
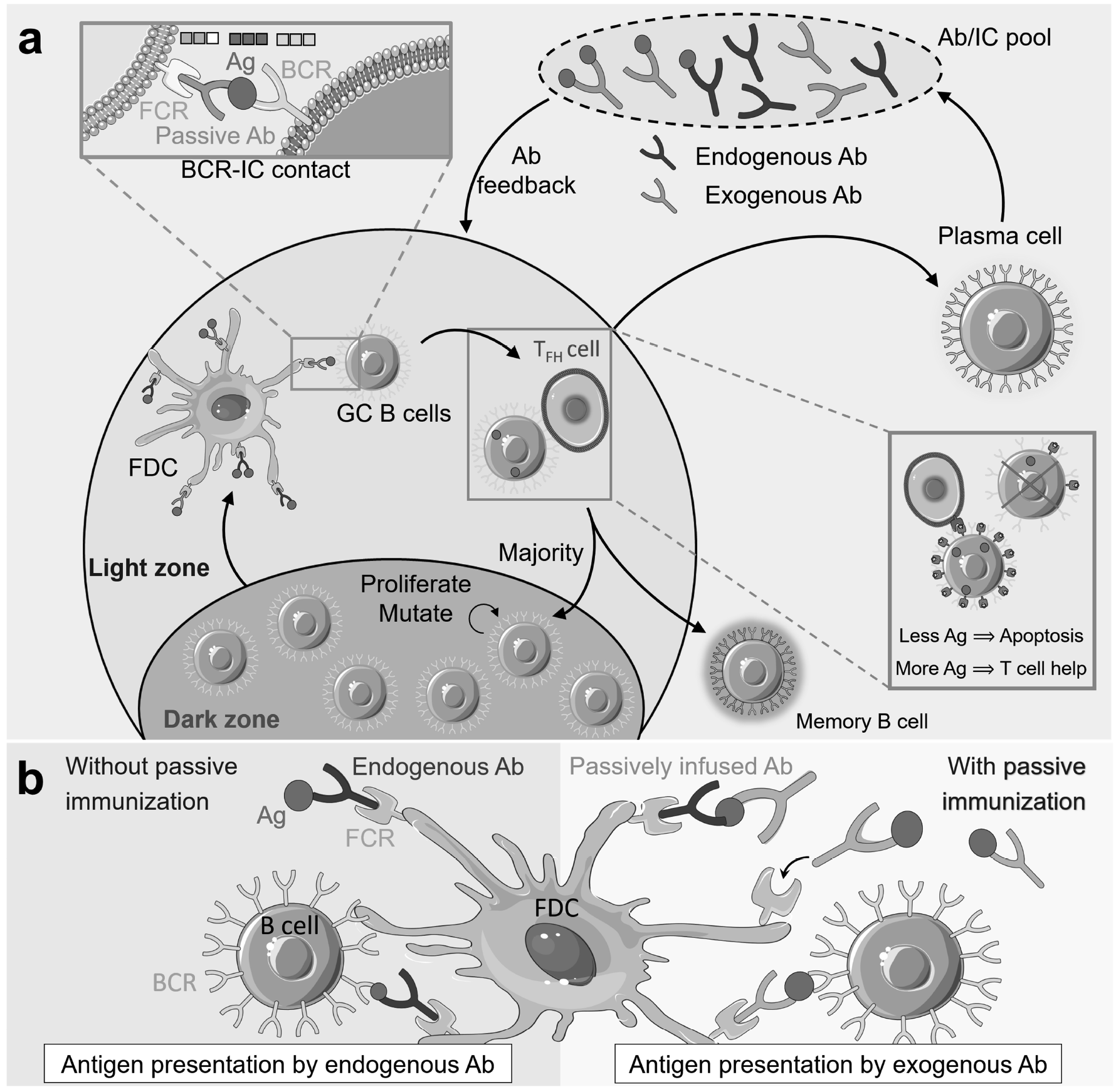
Schematic of the mechanism with which exogenous Abs influence the GCR. **a**, Schematic of the GCR: GC B cells in the light zone interact with Ag-presenting FDCs, forming BCR-IC contacts (left zoom). BCRs with greater affinity for Ag than the complexed Ab rupture the IC and internalize Ag. Acquisition of more Ag results in an increased probability of binding with T_*fh*_ cells and survival (right zoom). A majority of the selected B cells migrate to the dark zone while the remaining differentiate into Ab-producing plasma cells and memory B cells. Endogenous Abs, produced by plasma cells, and infused exogenous Abs, diffuse into the GC and capture Ag from existing ICs to form new ICs, or form higher affinity ICs in the serum and enter the GC, altering the selection stringency for B cells. **b**, Schematic of B cell-FDC interaction. Without passive immunization (left), endogenous Abs present Ag to B cells as ICs on FDCs. With passive immunization (right), administered exogenous Abs with higher affinity for Ag either capture Ag from pre-existing lower affinity ICs or react with Ag molecules in serum to form new ICs that are trafficked on to FDCs. B cells must thus acquire Ag from these stronger ICs, increasing B cell selection stringency.

We recognize that Abs in circulation can enter GCs, a phenomenon termed Ab feedback.^14,18,19^ Administered, or exogenous, Abs too can thus enter GCs. When the latter Abs have a higher affinity for Ag than endogenous Abs, they can preferentially form ICs by binding free Ag or prising Ag away from endogenous ICs displayed on FDCs, and themselves then be displayed as ICs on FDCs (Figure 1*b*). Indeed, ICs with higher Ab-Ag affinity have been shown to replace ICs with lower Ab-Ag affinity on FDCs.^14,18,19^ BCRs would then have to acquire Ag complexed with the higher affinity exogenous Abs, thus raising the selection stringency in the GC, potentially expediting and enhancing AM and modulating the humoral response. Tuning the affinity of the administered Ab for Ag would then be a handle to engineer the humoral response.

To test this hypothesis, we constructed an *in silico* model of the GC reaction (GCR) in the presence of passively administered Abs and performed stochastic simulations.

## Results

### Stochastic simulation model of the GCR and the influence of passive immunization

We considered the GCR in response to a single, non-mutating epitope, akin to simple immunogens such as haptens employed in numerous experiments^14,23–30^. We modelled the GCR using spatially homogeneous, discrete generation Wright-Fisher simulations (Figure 1*a*; Methods). We initiated the simulations with *N* = 1000 GC B cells expressing BCRs with low affinity for the Ag. We represented BCR paratopes and the antigenic epitope as bit-strings of length *L* with an alphabet of size 4. In each generation, also called a GC cycle, each B cell interacted on average with *η* ICs on FDCs. *η* was thus a surrogate for Ag availability in the GC. The affinity of a BCR for Ag was set proportional to the match length, *ε*, which we defined as the length of the longest common sub-string between the two sequences. In each B cell-IC encounter, the probability that the B cell acquired Ag was thus proportional to *ε* − *ω*, where *ω* was the match length of the Ag with the Ab in the IC. Following Ag acquisition, B cells competed for T_*fh*_ help. The probability that a B cell received T_*fh*_ help was proportional to the relative amount of Ag it acquired; the probability was 0 for a B cell that acquired no Ag and rose to 1 for a B cell that acquired the maximum Ag in the cycle. Accordingly, B cells with high amounts of internalized Ag relative to their peers were preferentially selected. Of the selected B cells, which we restricted to a maximum of 250, 10% differentiated equally into plasma and memory B cells, and 90% migrated to the dark zone, where they proliferated and mutated. Each B cell proliferated twice on average in one cycle. We let 10% of the daughter cells harbour a single, random mutation in their BCR sequences. The B cells then returned to the light zone and formed the pool of cells for the next cycle.

As AM progresses, lower affinity ICs on FDCs are continuously replaced by higher affinity ICs via Ab feedback^14,18^ (Figure 1*b*). We modelled this IC turnover as follows. During “natural AM”, *i.e.,* in the absence of passive immunization, we chose plasma cells from a previous generation randomly and let Abs produced by them form the ICs for the present generation. With passive immunization, because IC turnover is fast^14^, until endogenous Abs of higher affinity than the administered Abs for Ag are formed, ICs are expected predominantly to contain administered Abs. We therefore let, as an approximation, all ICs contain administered Abs, which we assumed were not limiting, until the average BCR affinity in a cycle exceeded the affinity of the administered Abs for the Ag, at which point, we switched to natural AM.

The GCR ended when B cells with BCRs of the highest match length (*ε* = *L*) were produced, marking completion of AM, or if all the B cells in a cycle died, representing GC collapse.

### Model recapitulates AM without passive immunization

We first examined whether the model recapitulated known features of the GCR in the absence of passive immunization. We set *L* = 3 for ease of computation and so that the BCR/Ab paratopes could be classified as having low (*ε* = 1), intermediate (*ε* = 2) or high affinity (*ε* = 3) for the Ag. We initiated the GCR with a random mixture of all types of B cells with low affinity for Ag. Note that there are 27 types of low affinity B cells, given the 3 different mutations possible from the defined Ag sequence for an alphabet of size 4 at each of the *L* = 3 positions. We performed simulations mimicking scenarios where Ag availability was not low (*η* = 8; see below).

Upon initiation of the GCR, the low affinity GC B cell counts dropped and gave way to the fitter, intermediate affinity B cells (Figure 2*a*). The intermediate affinity B cells rose in number, but soon let mutations give rise to the even fitter, high affinity B cells, which eventually dominated the GC B cell population. The average affinity of the B cell population for Ag increased gradually, showing AM (Figure 2*b*). Concomitantly, the average affinity of the Abs in the ICs presented on the FDCs increased, indicating increasing selection stringency (Figure 2*b*). The plasma cell output, which included B cells of intermediate and high affinity for Ag, also increased and saturated (Figure 2*c* inset). For the parameters chosen, AM reached completion in 2 – 3 weeks, in agreement with the observed temporal evolution of Ab affinity for haptens during natural AM^14,27,29,31^. The humoral response mounted by the GC, characterized by the instantaneous and cumulative affinity-weighted Ab output from the GC (Methods), intensified with time (Figure 2*c*). The long plasma cell half-life (~ 23 days^32,33^) implied that the humoral response would take longer to saturate than the duration studied in our simulations. Our focus was on AM; we therefore ignored processes beyond the completion of AM, including the eventual extinguishing of GCs due to Ag decay^34^. The above predictions matched previously observed characteristics of natural am^14,30,34,35^, indicating that the essential features of the GCR were captured by our simulations.

**Figure 2.**
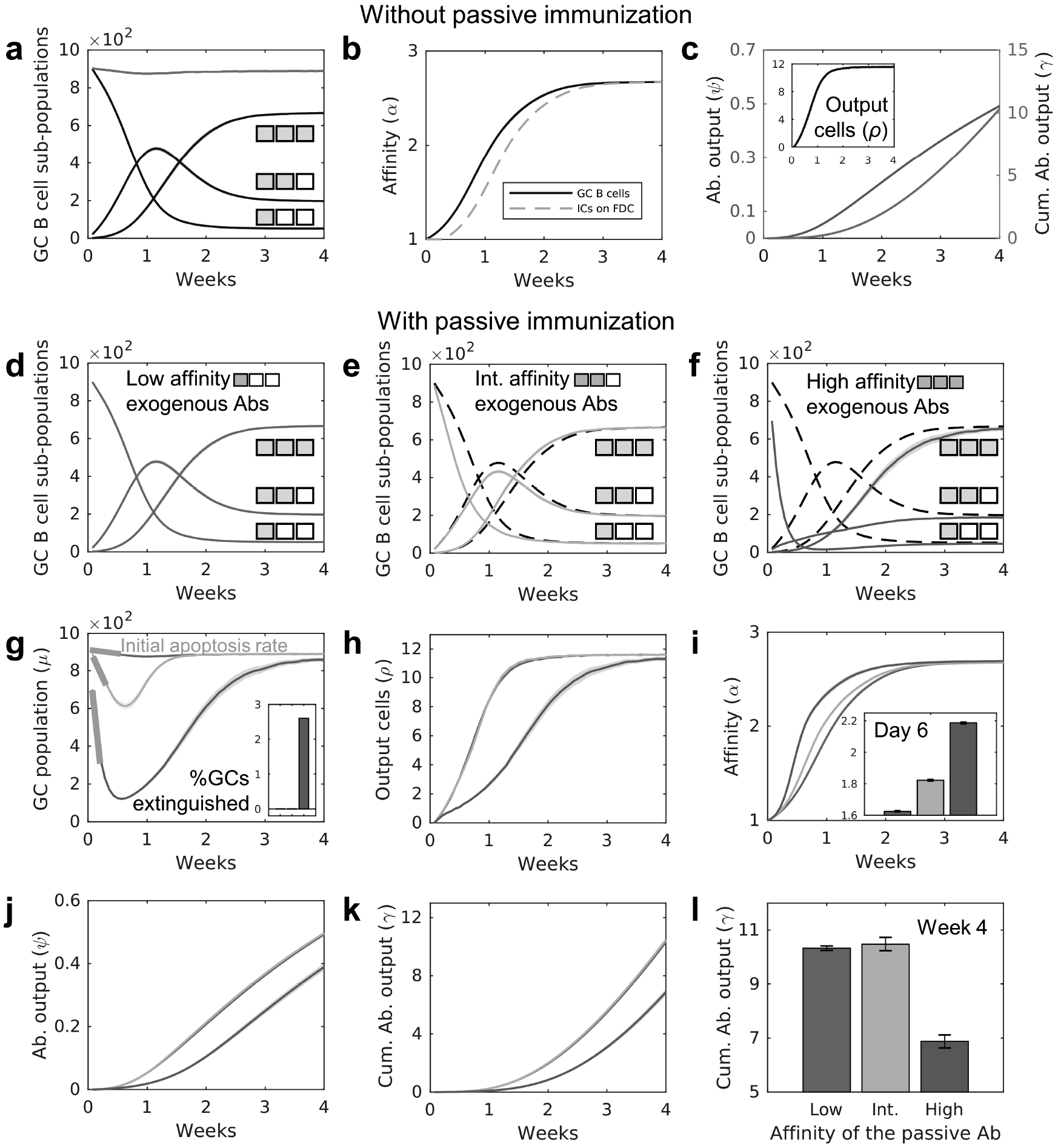
Passive immunization alters affinity maturation and the quality-quantity trade-off constraining the GCR. **a-c**, Natural AM. Temporal evolution of (a) low (*ε* = 1), intermediate (*ε* = 2) and high (*ε* = 3) affinity GC B cell sub-populations, and overall GC size, *μ*(*t*) (red), (b) the average affinity of GC B cells (black) and of the Abs in ICs (dashed orange), (c) the plasma cell output (inset), instantaneous Ab production from GCs, and the cumulative Ab output from GCs. The match length, *ε*, is illustrated with coloured positions in box-strings. **d-i** AM with passive immunization. Evolution of GC B cell sub-populations upon passive immunization with Abs of (d) low (*ω* = 1, grey), (e) intermediate (*ω* = 2, yellow) or (f) high (*ω* = 3, green) affinity. Dashed lines represent natural AM, shown for comparison. The match lengths are illustrated with green (endogenous) or blue (exogenous) coloured positions in box-strings. Temporal evolution of GC attributes upon immunization with low (grey), intermediate (yellow) and high (green) affinity Abs: (g) size (*μ*(*t*); % GCs extinguished over the course of 4 weeks shown in inset and initial apoptosis rates in blue), (h) plasma cell output (*ρ*(*t*)), (i) affinity (*α*(*t*)), (j) instantaneous Ab output (*ψ*(*t*)), and (k) overall Ab output (*γ*(*t*)). (l) Comparison of *γ* at week 4 shows that intermediate affinity Abs maximized the quality-quantity trade-off.

### AM with passive immunization reveals a quality-quantity trade-off constraining the GCR

We examined next how passive immunization altered the GCR. We performed simulations with passively administered Abs of low (*ω* = 1), intermediate (*ω* = 2) or high (*ω* = 3) affinity for Ag, which we assumed were available from the start of the GCR. With low affinity passive Abs, *i.e.,* when *ω* = 1, no IC turnover with exogenous ICs was possible. The GCR was thus indistinguishable from natural AM (Figure 2*d*). With *ω* = 2, AM proceeded faster than natural AM. Low affinity B cell counts declined faster than with natural AM because of the greater selection stringency imposed by the administered Abs (Figure 2*e*). The decline was even sharper with *ω* = 3 (Figure 2*f*). Correspondingly, the overall GC B cell population initially declined, with the decline steeper with higher *ω*. With *ω* = 2, the population was rescued with the emergence of B cells with intermediate affinity BCRs (*ε* = 2). Although such BCRs did not have a fitness advantage over the passive Abs, they had a 50% chance of prising Ag away in an encounter with an IC. Such B cells therefore had a significant chance of receiving the requisite survival signals. Their populations thus rose. The lower population of low affinity B cells present compared to natural AM implied that fewer intermediate affinity B cells could be produced, resulting in the net fewer intermediate affinity B cells when *ω* = 2 than natural AM (Figure 2*e*). The intermediate affinity B cells gave rise to high affinity B cells, which eventually dominated the B cell population (Figure 2*e*). With *ω* = 3, however, the intermediate affinity B cells too had a strong selective disadvantage imposed by the higher affinity ICs. The intermediate affinity B cell population thus rose at a very low rate, based on chance events where BCRs acquired Ag from ICs with a probability far smaller than 50% (Figure 2*f*). Mutations eventually yielded BCRs with high affinity, following which the GC B cell population was rescued and AM reached completion.

These results revealed a fundamental constraint on the GC response. As the selection stringency (*ω*) increased, the GCs experienced greater apoptosis of the lower affinity B cells (Figure 2*d-g*). Indeed, with *ω* = 3, a few GCs were extinguished (Figure 2*g* inset). The overall plasma cell output thus suffered as *ω* increased (Figure 2*h*). However, the GC B cells selected were of higher affinity because of the higher selection stringency (Figure 2*i*). The difference was evident from the average affinity of the GC B cells at day 6 (Figure 2*i* inset). Thus, as selection stringency increased, the average affinity for Ag, or quality, of the GC B cells selected increased, but the number, or quantity, of the GC B cells selected suffered. This quality-quantity trade-off implied that intermediate selection stringency maximized the GC response. The instantaneous and cumulative Ab outputs were thus highest with *ω* = 2 (Figure 2*j-l*). (The small *L* makes it difficult to distinguish between the cumulative outputs with low and intermediate *ω*; the difference is evident with *L* = 8 (Supplementary Figure S1).)

### Simulations capture experimental observations of the quality-quantity trade-off

Recent experiments present evidence of the quality-quantity trade-off constraining the GC response. In primed mice, passively administered Abs of high affinity for the Ag at the start of the GCR enhanced AM but also resulted in amplified GC B cell apoptosis and lower plasma cell output, whereas low affinity exogenous Abs maximized the GC B cell population and plasma cell output but resulted in relatively poor AM.^14^ Further, the effects of passive immunization were lost when the immunization was delayed. To test whether our model was consistent with these observations, we performed two sets of simulations: (i) GCs were started with low, intermediate or high affinity exogenous ICs to mimic early passive immunization, and (ii) GCs were allowed to undergo natural AM for 2 weeks, and then the ICs were turned over with administered Abs of low, intermediate or high affinity, to mimic delayed passive immunization. GC responses were calculated 2 and 4 days after passive immunization, following the experimental protocol^14^.

We found that early (week 0) passive immunization decreased the GC B cell population, and hence GC size, as the affinity of the administered Abs for Ag increased (Figure 3*a*). The decrease was due to increased GC B cell apoptosis (Figure 3*b*), which in turn resulted in decreased plasma cell output (Figure 3*c*). The affinity of the GC B cells, however, increased (Figure 3*d*), consistent with the quality-quantity trade-off. Late passive immunization, at week 2 from the start of the GCR, resulted in negligible differences in GC attributes (Figure 3*e-h*). By week 2, natural AM had nearly reached completion, so that endogenous Abs had affinities on par with if not higher than the administered Abs. The latter thus could not induce IC turnover and alter selection stringency. Our simulations, thus, reproduced qualitatively all the above observations^14^, reaffirming the quality-quantity trade-off constraining the GC response.

**Figure 3.**
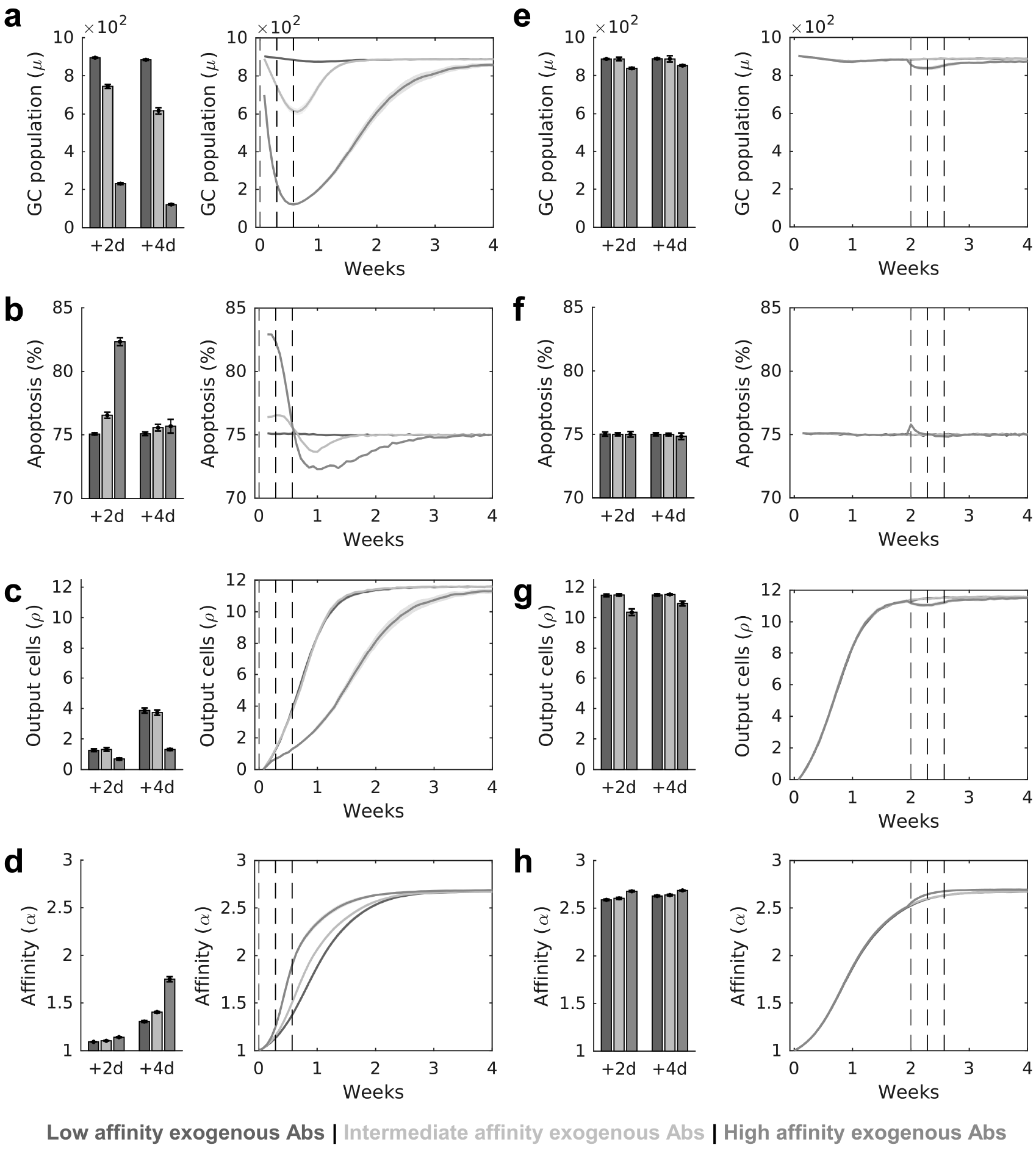
Simulations recapitulate experiments of the quality-quantity trade-off. Point measurements (left; 2 and 4 days after passive immunization) and temporal evolution of GC attributes (right), *viz.*, size, extent of apoptosis, plasma cell output and affinity after **a-d**, immunization with GC initiation, and **e-h**, immunization 2 weeks after GC initiation. Red and grey vertical dashed lines in all the temporal trends indicate the times of passive immunization and measurement of GC characteristics, respectively. The results are consistent with recent experimental observations^14^.

### Ag availability tunes the quality-quantity trade-off

In the simulations above, as the selection stringency increased, more B cell apoptosis occurred because B cells with low affinity BCRs failed to prise Ag away from ICs of higher affinity passive Abs. Each B cell had *η* attempts, on average, to acquire Ag from FDCs. *η* was a surrogate for the amount of Ag available, assuming all other factors were not limiting. We reasoned that if more Ag were available, leading to higher *η*, each B cell would have more chances to acquire Ag and therefore survive. The probability of Ag acquisition per attempt would remain unaltered, still favouring higher affinity B cells. Thus, higher GC output quantities could be realized without compromising quality.

To test this hypothesis, we performed simulations with low (*η* = 6) and high (*η* = 10) Ag availability following passive immunization with Abs of intermediate and high affinity for Ag (Figure 4). We found that higher *η* resulted in higher GC B cell populations, indicating greater B cell survival (Figure 4*a*). This in turn resulted in increased plasma cell output (Figure 4*b*). Importantly, the mean affinity of the GC B cells was unaffected by *η*, indicating that AM and the quality of the GC response was not affected by Ag availability (Figure 4*c*). AM was influenced by the affinity of the passively administered Abs, in keeping with our results above (Figures 2 and 3). To emphasize our claims, we examined the cumulative Ab output at week 4 (Figure 4*d*). The GC output was reduced by higher affinity passive Abs but was restored by increasing Ag levels: The high Ag regime resulted in ~ 84% and ~ 243% higher Ab outputs than the low Ag regime upon passive immunization with intermediate and high affinity Abs, respectively (Figure 4*d*). Ag availability thus tuned the quality-quantity trade-off; increased Ag availability offset the adverse effects of selection stringency on the quantity of the GC output without compromising its quality.

**Figure 4.**
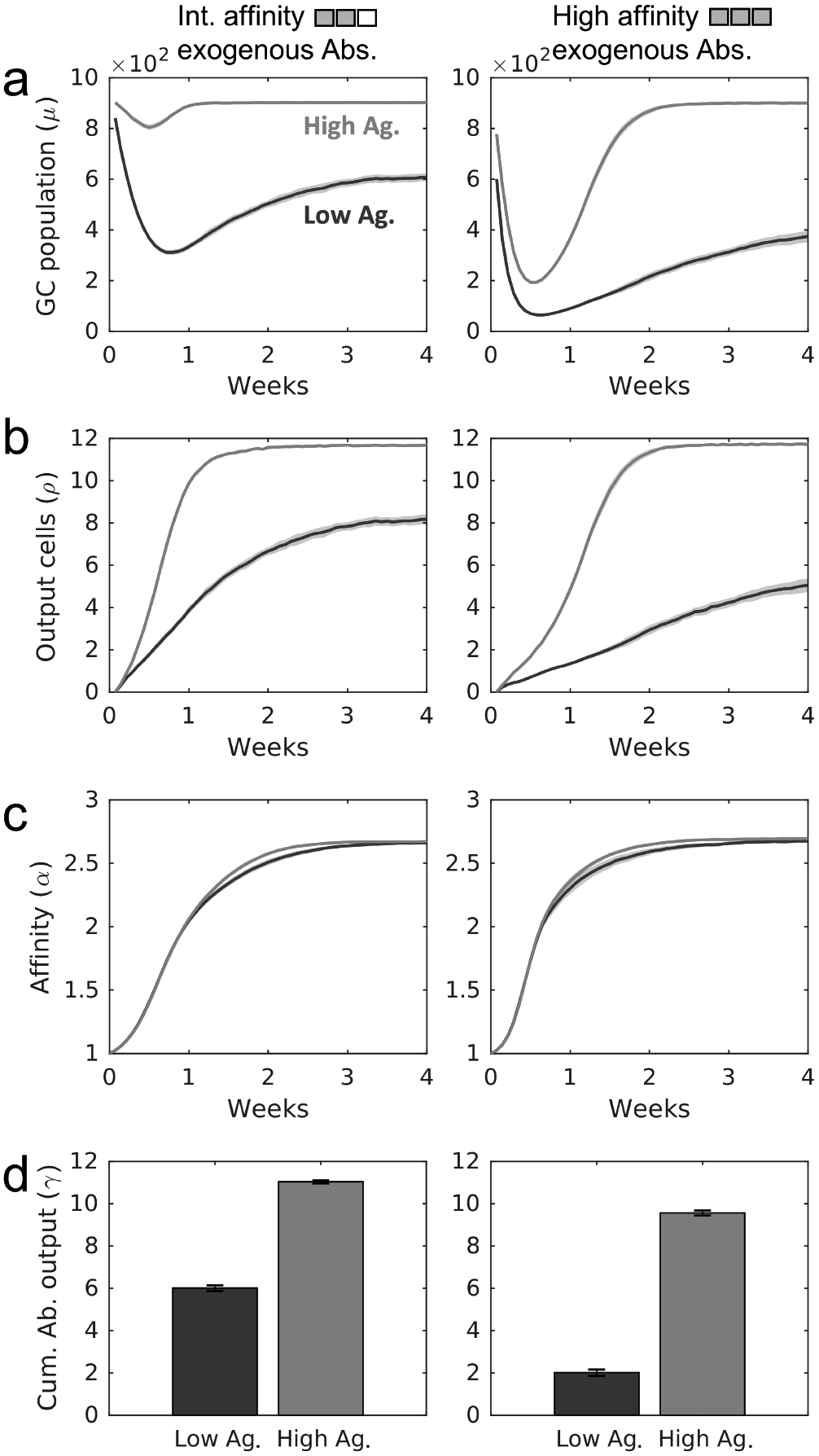
Ag availability tunes the quality-quantity trade-off in the GCR. GC characteristics: **a**, size, **b**, plasma cell output, **c**, affinity of GC B cells, and **d**, cumulative Ab output, at week 4 upon passive administration of exogenous Abs with intermediate (left column) and high (right column) affinity for Ag under low (*η* = 6, blue) and high (*η* = 10, magenta) Ag availability regimes.

### Simulations recapitulate experimental observations of the GC response with varying Ag availability

The above simulations were performed by keeping *η* constant in each simulation, thus representing a scenario where the Ag level did not vary during the GCR. Ag levels could, however, vary during the GCR. In a recent study, Ag availability was varied by administering varying doses of Ag during the GCR in the absence of passive immunization. ^33^ Interestingly, an exponentially increasing Ag dosage, mimicking the scenario where Ag levels rise due to pathogen replication following an infection, was found to induce better GC responses than scenarios where the dosage was constant or fell with time.^33^ We examined whether our simulations could recapitulate the latter observations.

In the experiments^33^, Ag availability was modulated by varying the administered Ag doses in an exponentially-increasing, constant or exponentially-decreasing manner while maintaining constant overall amounts of infused Ag. To mimic these scenarios, we performed simulations of natural AM with *η* varying within each simulation following the patterns of the dosages employed, but with a constant area under the curves until day 7, when Ag loading ended (Figure 5*a*). *η* was subsequently assumed to decay exponentially at a rate commensurate with IC turnover on FDCs.

**Figure 5.**
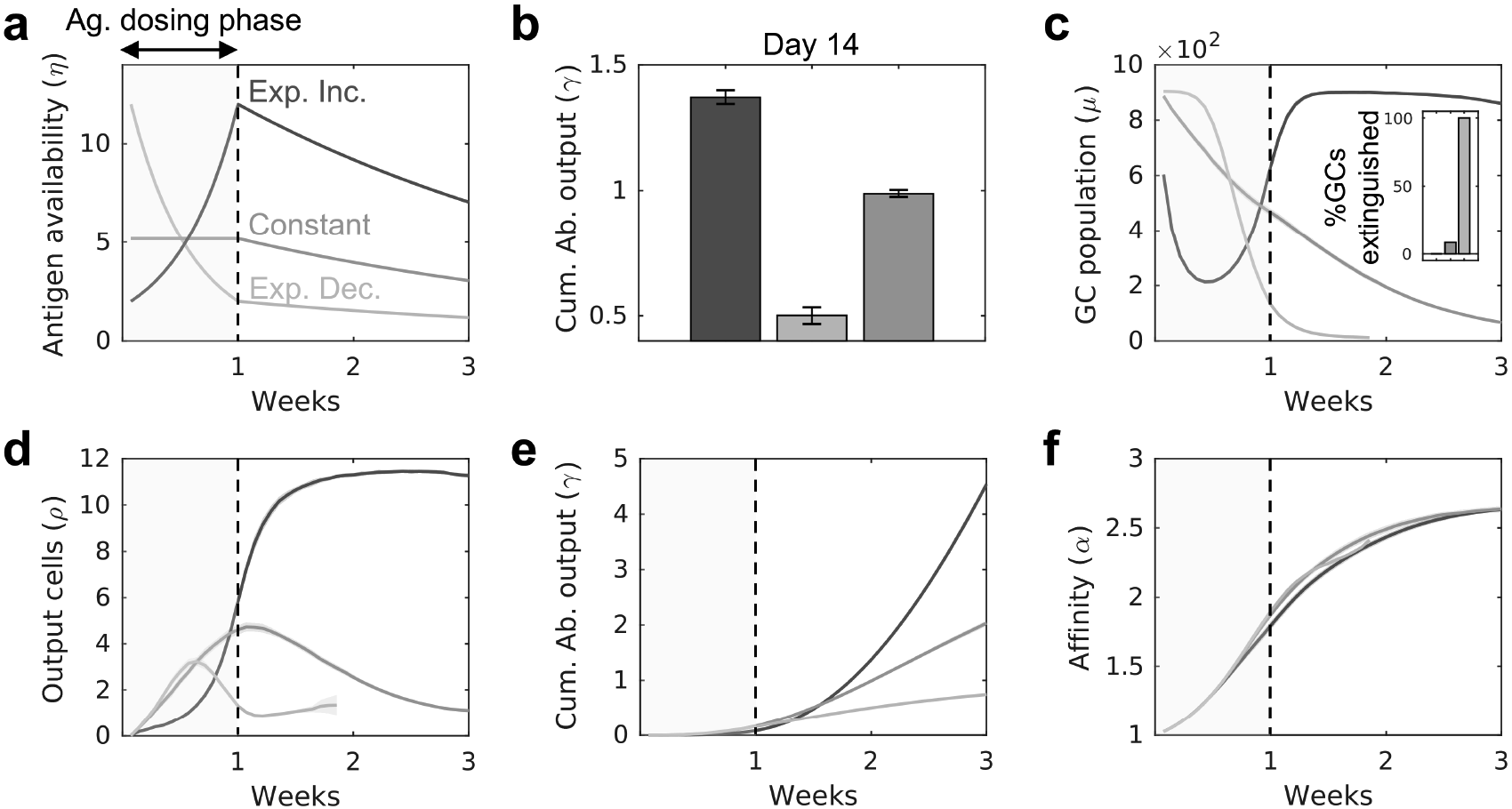
Simulations recapitulate experiments with varying Ag availability. **a**, Time evolution of Ag availability, *η* (see text), mimicking the exponentially-increasing, constant or exponentially-decreasing Ag dosage till day 7 (Ag dosing phase), and exponential decline afterwards. The latter decline occurs with the rate constant of 0.018 generation^−1^. (ICs on FDCs are lost by turnover with an estimated rate constant^32,33^ of 0.012 generation^−1^. We let an additional 50% loss occur, as an approximation, due to Ag acquisition by B cells.) **b**, GC responses elicited by the three Ag dosing profiles in terms of the cumulative Ab output on day 14. **c-f**, Temporal evolution of the GC size (% GCs extinguished by week 3 in inset), plasma cell output, cumulative Ab output, and GC B cell affinity, respectively. The results are consistent with corresponding experiments^33^.

We found that the exponentially increasing *η*, which ensured adequate Ag availability in GCs, yielded the highest cumulative Ab output on day 14 (Figure 5*b*), in agreement with experiments^33^. Constant and exponentially decaying Ag dosing protocols led to progressive apoptosis, GC collapse and abrogation of plasma cell output by week 3, while exponentially increasing Ag ensured GC survival and robust plasma cell outputs (Figure 5*c, d*). The exponentially increasing dosing also yielded the highest overall Ab output (Figure 5*e*). Yet, no significant differences in AM were observed between the different Ag dosing profiles (Figure 5*f*), in keeping with our simulations above of the decoupling between the quality and the quantity of the GC response with high Ag availability (Figure 4). The observed dependence of the GC output on the Ag loading profile was consistent with experimental observations^33^.

Our simulations thus recapitulated and synthesized independent experiments^14,33^, giving us confidence that our formalism accurately captured the essential features of the GCR in the presence of passively administered Abs. We applied our simulations next to identify optimal passive immunization protocols.

### *In silico* identification of optimal sequential passive immunization protocols

Our results above indicate that the affinity of passively administered Abs as well as Ag availability define the GC response. Ag availability is expected to be determined by the pathogen load in infected individuals. We therefore asked whether passive immunization protocols could be identified that would maximize the GC response for given Ag availability. We assumed, following recent experiments^14^, that Abs with low, intermediate, and high affinity for the target Ag were available. These could then be administered in different sequences. Following recent protocols of passive immunization with HIV-1 bNAbs^36^, we examined a strategy where 3 Ab infusions could be administered. For simplicity, we assumed that the doses were equally spaced and let them be administered on day 0 (start of the GCR), day 3.5 (cycle 7) and day 7 (cycle 14) of the GCR (Figure 6*a*). (A cycle typically lasts 12 h.^16,28^) Each dose could be of low (denoted as “L”), intermediate (denoted as “I”), or high (denoted as “H”) affinity Ab. For example, the combination H-I-L referred to the infusion of high affinity Abs at cycle 0, intermediate affinity Abs at cycle 7, and low affinity Abs at cycle 14. The 3-dose protocol allowed 27 unique passive Ab administration sequences, from the all low, L-L-L, to the all high, H-H-H, affinity sequence. We performed simulations over a wide range of (constant) Ag availability, from *η* = 1 to *η* = 20, exploring each of the 27 Ab sequences for every integral *η* in the range. We calculated the weighted cumulative Ab output at day 14 (cycle 28) as a measure of the humoral response.

**Figure 6.**
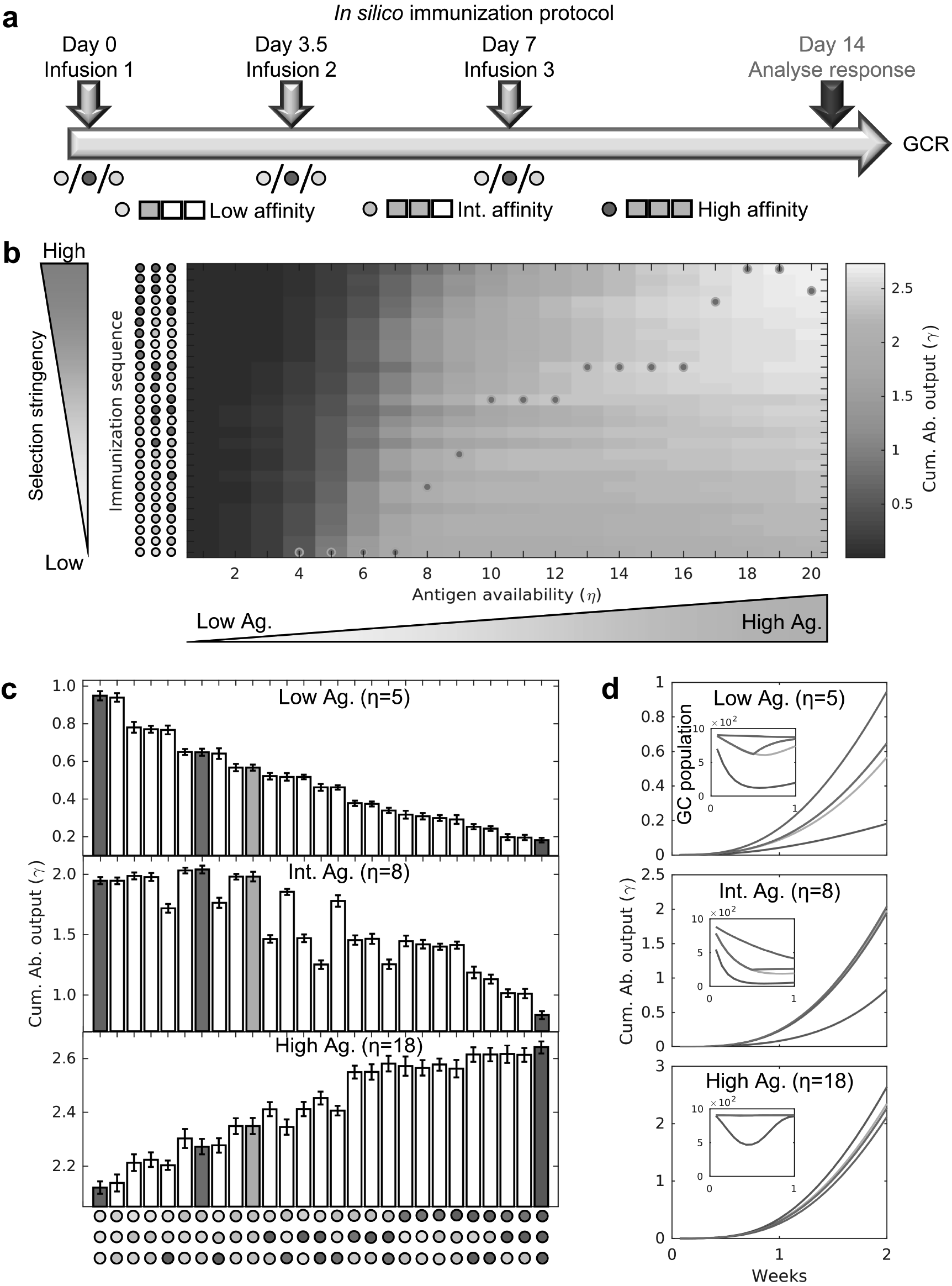
Optimization of Ab combinations for sequential passive immunization. **a**, Schematic of the *in silico* passive immunization protocol, with 3 doses of either low (grey dot), intermediate (yellow dot) or high (green dot) affinity Abs on days 0, 3.5 and 7 after initiation of the GCR, and measurement of response on day 14. **b**, The cumulative Ab outputs at day 14 elicited by all possible passive Ab administration sequences as functions of Ag availability, *η*. **c**, Cumulative Ab outputs for all possible passive Ab administration sequences for low (*η* = 5), intermediate (*η* = 8) and high (*η* = 18) Ag availability. The responses to the sequences L-L-L, I-I-I, H-H-H, and I-L-I (see text) are coloured grey, yellow, green, and red, respectively. **d**, Temporal evolution of the cumulative Ab output (see Supplementary Figure S2 for evolution of plasma cells and affinity) and GC size (inset) for the latter sequences.

The humoral responses elicited are summarized as a heat-map (Figure 6*b*). The Ab sequence eliciting the best response for each *η* is indicated by a red dot. For example, the best sequences for *η* = 5 (low), 8 (intermediate), and 18 (high) were L-L-L, I-L-I and H-H-H, respectively. To understand these optimal combinations, we examined the humoral responses to each of the 27 sequences for these values of *η* (Figure 6*c*). Selection stringency increased as the usage of high affinity Abs increased in the sequences. We found, interestingly, that the response gradually weakened as the selection stringency increased for *η* = 5. With *η* = 18, the opposite happened, with the response the best when selection stringency was the highest. With *η* = 8, an intermediate level of selection stringency elicited the best response. These results were in keeping with our observations above where increasing Ag availability relaxed the constraint imposed by the quality-quantity trade-off, allowing higher quantity and quality of the GC response with higher selection stringency. To elucidate these findings further, we examined the dynamics of the GCR for four different dosing sequences, L-L-L (gray), I-I-I (yellow), H-H-H (green) and I-L-I (red), for the three values of *η* above (Figure 6*d* and Supplementary Figure S2). With *η* = 5, L-L-L resulted in a low selection threshold and maintained the plasma cell output, while the harsher selection imposed by higher affinity passive Abs in the other combinations led to reduced Ab output, GC size, and plasma cell output (Figure 6*d* and Supplementary Figure S2). With *η* = 8 and *η* = 18, however, the GCs withstood higher selection stringencies, yielding better outputs with immunization sequences containing higher affinity Abs. In all cases, the quality of the GC response, defined by the average GC B cell affinity, increased with the usage of higher affinity passive Abs (H-H-H > I-I-I > I-L-I > L-L-L; Supplementary Figure S2).

In summary, for a given Ag availability, our simulations enabled the identification of the sequence of administration of Abs that maximized the GC response. Broadly, sequences containing lower affinity Abs were optimal with low Ag availability, whereas sequences containing higher affinity Abs elicited the best response with high Ag availability.

## Discussion

With growing evidence of its ability to elicit improved and lasting humoral responses, passive immunization is poised to transcend its classical role as a strategy for the temporary alleviation of disease burden and emerge as a potent tool to engineer host humoral responses. Here, we elucidated the mechanism with which administered Abs can influence endogenous Ab production, addressing a fundamental knowledge gap that has precluded the rational deployment of passive immunization to engineer humoral responses. We hypothesized that administered Abs raise the selection stringency in GCs by preferentially forming ICs and preventing B cells with low affinities for Ag from acquiring Ag and receiving survival cues. Stochastic simulations of the GC reaction based on this hypothesis captured several independent experimental observations. The simulations unravelled a quality-quantity trade-off constraining the GC reaction and showed how optimal passive immunization protocols can be designed to exploit this trade-off and maximize the GC output.

Previous mathematical models and stochastic simulations of AM have presented important insights into the GCR, such as the cycling of B cells between the light and dark zones^21,32^ and the limiting nature of T_*fh*_ help^37,38^, and have suggested vaccination protocols that could guide the GCR into eliciting desired Ab responses^33,34,39,40^. Models have also explored the evolution of B cells yielding bNAbs of HIV-1 and the reasons impeding their selection.^41–43^ The influence of passive immunization has been less well explored. One study argued that passive immunization increases the selection stringency in GCs via epitope masking, where administered Abs bind Ag in ICs already presented on FDCs and block BCRs from accessing Ag.^14^ The resulting model made predictions consistent with the quality-quantity trade-off. The model, however, assumed the binding of administered Abs to the ICs to be reversible, but did not account for the possibility that Ag could be tugged away by the dissociating Abs based on the relative affinities of the endogenous and exogenous Abs for Ag. Our model accounted for the latter possibility and explained not only the quality-quantity trade-off, but also the observed IC turnover and the long residence times on the FDC surface of high but not low affinity passive Abs^14^. A new, more robust conceptual view of the selection forces driving AM in GCs and the influence passive immunization has on these forces thus emerges.

Our findings synthesize several independent experimental observations. By showing how AM is influenced not only by the affinity of the administered Abs for Ag but also by Ag availability, our findings recapitulated observations of the GC response with passive immunization using Abs of different affinities^14^ as well as with active immunization using Ag dosing protocols that tuned Ag availabity^33^. Our findings are also consistent with improvements in the endogenous Ab response elicited by passive immunization in several studies. For instance, in the newborn or 1 month old rhesus macaque model of SHIV infection, administration of neutralizing Abs enhanced endogenous neutralizing Ab production, which suppressed set-point viremia, delayed disease progression and enhanced survival.^12,13^ Similarly, passive immunization of mice with anti-CD4 binding site Abs enhanced the production of Abs targeting the CD4 binding site of HIV-1 gp120.^11^ More recently, improved neutralization breadth and potency of endogenous Abs was observed following passive immunization of HIV-1 infected individuals with the HIV-1 bNAb 3BNC117.^15^ Further, in the latter study, better responses were elicited in individuals not on antiretroviral therapy than in those on antiretroviral therapy. Antiretroviral therapy suppressed viremia. The former individuals thus had higher plasma viremia than the latter, which could lead to greater Ag availability in the GCs and therefore better GC responses, consistent with our predictions. The affinity of administered Abs for Ag and the availability of Ag thus emerge as potentially tunable handles to engineer the host humoral response using passive immunization strategies.

Although Ag availability within GCs remains difficult to estimate^33^, it is likely to be proportional to the pathogen load. Pathogen loads can vary widely across individuals; for instance, the set-point viral load varies by 4 orders of magnitude across HIV-1 infected individuals^44^. Our simulations suggest that passive immunization protocols must be personalized based on the pathogen load. By comprehensively spanning parameter space, the simulations showed that passive immunization with Abs of low affinities for Ag were optimal when Ag availability in the GCs was low, whereas higher affinity Abs were best suited when Ag availability was high. Future studies may also examine whether simultaneously tuning Ag availability, using suitable Ag administration patterns^33^, and selection stringency in the GCs, using passive immunization with Abs of suitable affinities, may be a more powerful strategy than passive immunization alone. Our simulations present a framework for the design of such optimal passive immunization protocols. Finally, because passively administered Abs alter AM by forming ICs in our simulations, our simulations can also be applied to define optimal strategies of passive immunization with ICs, which have been argued to improve the humoral response more than with the uncomplexed Abs^45–47^.

Our simulations have been restricted to describing the GCR with a single, non-mutating antigenic epitope. With rapidly evolving pathogens such as HIV-1, B cell evolution is expected to occur in the presence of multiple epitopes in a GC. Accounting for this diversity would be an important extension of our formalism, especially given that the strength of the endogenous response following passive immunization was found to correlate with antigenic diversity^15^. Further, our simulations did not include T cell receptor specificity that allows interactions of T_*fh*_ cells only with cognate GC B cells. Both multiple epitopes and T cell receptor specificity may be important to the development of breadth in Ab responses.^34,41,43^ We employed approximate descriptions of the fitness of B cells based on the energetics of BCR-IC interactions, following previous studies^41^. More detailed descriptions based on intracellular signalling following BCR stimulation, leading to B cell proliferation and differentiation or apoptosis, may yield more reliable fitness landscapes. Such landscapes would be important to understanding recent observations where stronger immune responses were elicited by upregulating the inhibitory receptors on B cells and raising the signalling threshold for B cell selection. ^48^

In summary, our simulations present a mechanistic understanding of the influence of passive immunization on the GCR, capture independent experimental observations, and facilitate optimization of passive immunization protocols to engineer the humoral response.

## Methods

### Stochastic simulation model of the GCR

#### Initialization

Low affinity seeder B cells initiate the GCR and proliferate rapidly until the GC population reaches a steady state of ~ 1000 cells, at which point somatic hypermutation of the immunoglobulin genes of the GC B cells switches on.^28,34,49,50^ We thus initiate our simulations with *N* = 1000 low affinity B cells. We consider AM of the B cells in response to a single, non-mutating antigenic epitope, representative of haptens or other simple Ags employed widely^14,23–30^. The Ag epitope and BCR/Ab paratopes are represented as bit-strings of length *L* and an alphabet of size 4, similar to previous AM models^34,41,51^. The size of the alphabet approximates the classification of amino acids into positively charged, negatively charged, polar and hydrophobic groups. ^41^ We first choose a randomly assembled string to represent the Ag. We then choose a BCR paratope for each cell in the initial B cell pool by randomly mutating the Ag sequence at *L* – 1 randomly chosen positions. We subject the B cell pool to selection, mutation, and proliferation in discrete generations, described below.

#### Antigen acquisition and T_fh_ cell help

Let *N*_*GC*_(*t*) be the GC B cell population in the *t*^*th*^ generation, or cycle. We let the B cells acquire Ag as follows. We select a B cell randomly and let it interact with an IC on the FDC, also chosen randomly. The ICs available on the FDCs are described below. The B cell succeeds in acquiring an Ag molecule during this encounter with the probability *f*_*Ag*_, defined below. This process is repeated *N*_*GC*_(*t*) × *η* times with replacement, so that each B cell has an average of n chances to acquire Ag in each generation. The B cells are then subjected to competition for *T*_*fh*_ cell help. A B cell is chosen at random for *T*_*fh*_ help, and allowed to survive with a probability 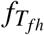, also defined below. This process is repeated *N*_*GC*_(*t*) times or until 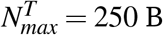 cells are selected, whichever is earlier.

#### Proliferation, evolution, and GC exit

Of the selected cells, we randomly select 90% for proliferation (in the dark zone) and let the remaining 10% differentiate equally into plasma and memory B cells and exit the GC^34^. Each B cell selected for proliferation proliferates twice on average.^24,34^ We then select 10% of the daughter cells and introduce single random point mutations in each of their BCR sequences. The resulting B cells form the pool for the next GC cycle.

#### IC turnover

FDCs mainly present ICs of Ag bound to Abs on FC*γ* or complement receptors.^18,19,52–54^ Previous studies have shown that Ag by itself may not bind with high affinity to Fc*γ* or complement receptors^55,56^ and therefore may not be efficiently presented on FDCs. Indeed, humoral responses have been found to be significantly better when ICs were used for passive immunization compared to uncomplexed Abs.^45–47^ Upon Ag stimulation, low affinity ICs are initially formed by endogenous broadly-reacting immunoglobulin M (IgM) Abs with Ag, which initiate the GCR.^33,57^ As the GCR proceeds, these low affinity ICs are replaced by higher affinity ICs by a process of Ab feedback. Endogenous Abs produced by plasma cells in the vicinity of the GC may re-enter the GC^14,18^ and capture Ag from existing ICs to form new ICs (Figure 1). Abs may also bind Ag in the serum and form ICs which are then deposited on the FDC via sub-capsular sinus macrophages and marginal zone B cells.^16,53,54,58,59^ Finally, higher affinity Abs may competitively displace low affinity Abs in existing serum ICs and be trafficked similarly on to FDCs. IC turnover from low to high affinities has been observed experimentally.^14^

We implement IC turnover as follows. For natural AM, we let affinity matured endogenous Abs secreted by the plasma cells in the *j*^*th*^ generation form the ICs presented on FDCs in the *j* + *τ*^*th*^ generation. In agreement with reported IC turnover time scales^14^, we let *τ* be 2 generations. AM is not sensitive to variations in *τ* (Supplementary Figure S3). With passive immunization, we let exogenous Abs form the ICs on FDCs until endogenous ICs of the same average affinity for Ag as exogenous Abs are formed, at which point we implement turnover akin to natural AM. It follows that higher affinity exogenous ICs show higher residence times on FDCs than lower affinity exogenous ICs, consistent with observations^14,18^.

#### B cell fitness for Ag acquisition

For each B cell, we define the match length, *ε*, as the length of the longest common sub-string of its BCR and Ag sequences^41,51^, and assume it to represent the strength of BCR-Ag binding. Similarly, the match length of the complexed Ab in the IC with Ag is *ω*. The fitness, *f*_*Ag*_, defined as the probability with which the B cell acquires Ag during a BCR-IC contact, depends on the relative magnitudes of *ε* and *ω* and can be derived from the free energy of spontaneous IC formation (Eq 1), spontaneous BCR-Ag complex formation (Eq 2) and competitive Ag acquisition by BCRs (Eq 3):

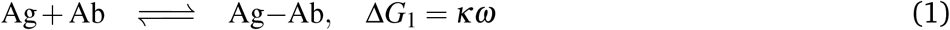

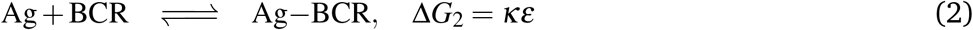

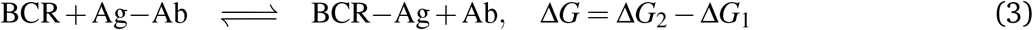

Here, *κ* is the constant, per-site free energy of binding of the Ab or BCR with the Ag. Similar to previous models^34,41^, steric factors and relative positions of the interacting epitopes in three-dimensional space are not considered. It follows that Δ*G* = *κ* (*ε* − *ω*). The relative fitness of a BCR is obtained by the normalization 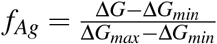, where Δ*G*_*min*_ = −*L*_*κ*_ represents the scenario where the Ab in the IC has the highest possible affinity for the Ag and the BCR has no affinity, and Δ*G*_*max*_ = *L*_*κ*_ represents the opposite. Simplifying, we obtain

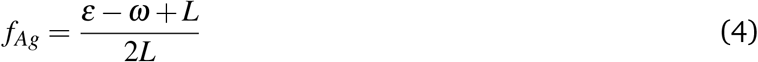

Other functional forms of *f*_*Ag*_ such as exponential landscapes have been considered (Supplementary Figure S4), and do not qualitatively change the model behaviour or predictions.

#### B cell fitness for T_*fh*_ cell help

B cells possessing relatively high amounts of internalized Ag, and therefore high pMHCII surface densities, compared to peers competitively elicit *T*_*fh*_ help.^16^ To decide whether a B cell succeeds, we let the amount of Ag acquired by the B cell be *θ*. (Note that this depends on *f*_*Ag*_ above.) Let *θ*_*min*_ and *θ*_*max*_ be the minimum and maximum amounts of Ag acquired across all B cells in the generation. The probability of the B cell successfully eliciting help from *T*_*fh*_ cells is then written as

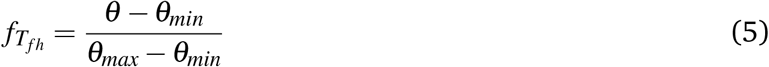

This description ensures that relative and not absolute amounts of acquired Ag determine *T*_*fh*_ help^16,25^, and the B cell internalizing maximum Ag always gets selected upon encountering a *T*_*fh*_ cell.

#### Termination

AM naturally ends when the highest match length (*ε* = *L*) B cells are produced. In realizations where the GC thrives, we terminate the GCR when *t*_*max*_ = 84 GC cycles is reached. In a fraction of the realizations, the GC B cell population goes to zero and the GC is extinguished. For every scenario, we perform 5000 realizations of the GCR to obtain reliable statistics.

### Measures of the GC response

We use the following measures to quantify the GC dynamics and response. To obtain standard errors of means, we divide the 5000 realizations in each scenario into 50 ensembles of 100 realizations each and obtain ensemble averages. The ensemble size is chosen to mimic a lymph node, which comprises approximately 100 GCs^33^.

#### Quality

The average instantaneous affinity of the GC B cells measures the quality of the GC response, and is defined as

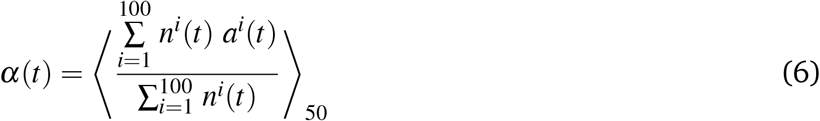

where *n*^*i*^(*t*) and *a*^*i*^(*t*) are the population and average affinity of the B cells in the *i*^*th*^ GC in an ensemble at GC cycle *t*, respectively. B cell affinity is defined in terms of the match length, *ε*. AM saturates when *α*(*t*) stabilizes at a time-invariant value.

#### Quantity

The average instantaneous GC size is defined as

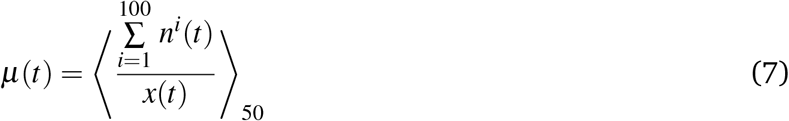

where *x*(*t*) is the number of surviving GCs in an ensemble at generation *t*. The quantity of the GC response is measured by the instantaneous output plasma cells per GC, *ρ*(*t*). We consider output plasma cells of *ε* ≥ 2 because only affinity matured GC B cells with BCRs that bind sufficiently strongly to Ag have been shown to differentiate into plasma cells^22^. Therefore,

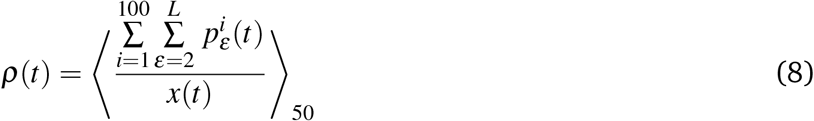

where 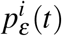 is the number of plasma cells with BCRs of match length ε output from the *i*^*th*^ GC in an ensemble at GC cycle *t*.

#### Quality and quantity combined

The effectiveness of the GC response is determined by a combination of serum Ab titres and their affinities for Ag. To estimate this effectiveness, we employ the following procedure. We define the instantaneous affinity-weighted plasma cell output of the GCs in a lymph node as

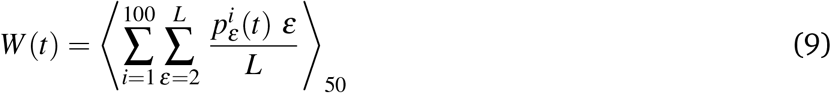

The affinity-dependent weights, 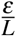, account for higher affinity plasma cells producing Abs that bind more efficiently to Ag and mount a stronger humoral response. Assuming that activated plasma cells die at the per capita rate *δ*_*p*_(= 0.015 per generation^32,33^), the affinity weighted cumulative plasma cell output at the *t*^*th*^ generation is

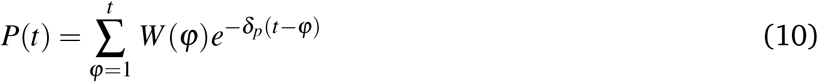

If each plasma cell produces Abs at the rate *β* = 8.64 × 10^7^ Abs per generation, which corresponds to the estimated 2000 Ab molecules/plasma cell/second^60^, the instantaneous affinity weighted Ab output at the *t*^*th*^ generation would be

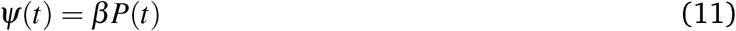

Abs are cleared from circulation at the per capita rate *δ*_*A*_(= 0.01165 per generation^32,33^). The cumulative size of the affinity weighted Ab pool at the *t*^*th*^ generation is therefore

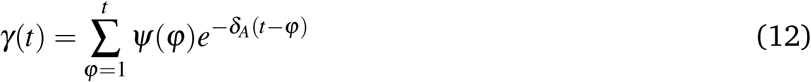

Thus, *α*(*t*) and *ρ*(t) are measures of the quality and quantity of the GC response, respectively, and *γ*(*t*) is a measure of the effectiveness of the response accounting for both quality and quantity. In the figures, *ψ*(*t*) and *γ*(*t*) are expressed in units of nmol Abs/generation/GC ensemble.

## Acknowledgements

We thank Rustom Antia for comments and the Wellcome Trust/DBT India Alliance Senior Fellowship IA/S/14/1/501307 (NMD) for funding.

## Author Contributions

A.K.G., R.D. and N.M.D. conceptualized and designed the study. A.K.G. and R.D. simulated the model. A.K.G., R.D. and N.M.D. analysed results and wrote the manuscript.

## Code availability

Codes employed in this study are available upon request.

